# A pancreatic cancer mouse model with human immunity

**DOI:** 10.1101/2023.05.24.542127

**Authors:** Norio Miyamura, Kodai Suzuki, Richard A. Friedman, Aristidis Floratos, Yuki Kunisada, Kazuya Masuda, Andrew M. Lowy, Moriya Tsuji, Kazuki N. Sugahara

## Abstract

Pancreatic ductal adenocarcinoma (PDAC) is characterized by a tumor immune microenvironment (TIME) that promotes resistance to immunotherapy. A preclinical model system that facilitates studies of the TIME and its impact on the responsiveness of human PDAC to immunotherapies remains an unmet need. We report a novel mouse model, which develops metastatic human PDAC that becomes infiltrated by human immune cells recapitulating the TIME of human PDAC. The model may serve as a versatile platform to study the nature of human PDAC TIME and its response to various treatments.

## Main

Immune checkpoint blockade (ICB) has become an important therapeutic option for multiple cancers including melanoma, non-small cell lung cancer, kidney cancer, esophageal cancer, and Hodgkin lymphoma^1, 2^. However, accumulating evidence suggests that many cancer types, such as PDAC, are resistant to ICBs, making it critical to understand the TIME in depth to develop effective immunomodulatory strategies that overcome resistance mechanisms^3–5^. Here, we developed a humanized PDAC (huPDAC) mouse model, which provides us with an important new tool to study the human PDAC TIME in mice. The model is comprised of orthotopic human PDAC in the presence of a full repertoire of functional human immune cells.

Mouse and human immunity have considerable differences, which presents challenges during the development and translation of immunotherapies for human diseases^6, 7^. Humanized mice, also known as human immune system (HIS) mice, in which human immune cells are engrafted into immunodeficient mice, originally emerged as a tool to study the pathogenesis of human immunodeficiency virus-type 1 (HIV-1)^8^. Since then, HIS mice have been used to study various human diseases including cancer^9–11^. However, it has been a major challenge to develop an ideal HIS mouse model, which carries a human leukocyte antigen (HLA) and has a high human CD45^+^ cell content with a full repertoire of functional immune cells^10^. As the platform for our huPDAC mice, we have used our original HIS mouse model, which is one of the few models that carry the desired features of an ideal HIS model listed above^12–19^. Our HIS mice are generated by engrafting cord-derived hematopoietic stem cells (HSCs) into immunodeficient NOD/SCID/IL2Rγ^-/-^ (NSG) mice that lack the mouse β2-microglobulin (Fig. 1a). Prior to the HSC infusion, genes encoding HLA-A*02 (HLA-A2) and human cytokines (huCytokines) that are essential for the hematopoiesis of human lymphoid cells, i.e., granulocyte macrophage colony-stimulating factor (GM-CSF), interleukin (IL)-3, and IL-15, are introduced into the mice by adeno-associated virus serotype 9 (AAV9)-mediated gene transfer, and mouse innate immune cells are ablated by a sub-lethal whole body irradiation. After 15 weeks, 80-95% of peripheral blood mononuclear cells (PBMCs) are reconstituted with human CD45 (hCD45)^+^ cells with equivalent proportions of T, B, and natural killer T (NKT) cells to those in humans. We have shown that the human CD8^+^ and CD4^+^ T cells, NKT cells, B cells, and dendritic cells (DCs) in the resulting HIS mice are fully mature and properly respond to various infections and vaccines^12–19^. Of note, the CD8^+^ T cells show a tumor antigen-specific response in an HLA-restricted manner making the HIS mice an optimal platform to study cancer immunity^14^.

**Figure 1.**
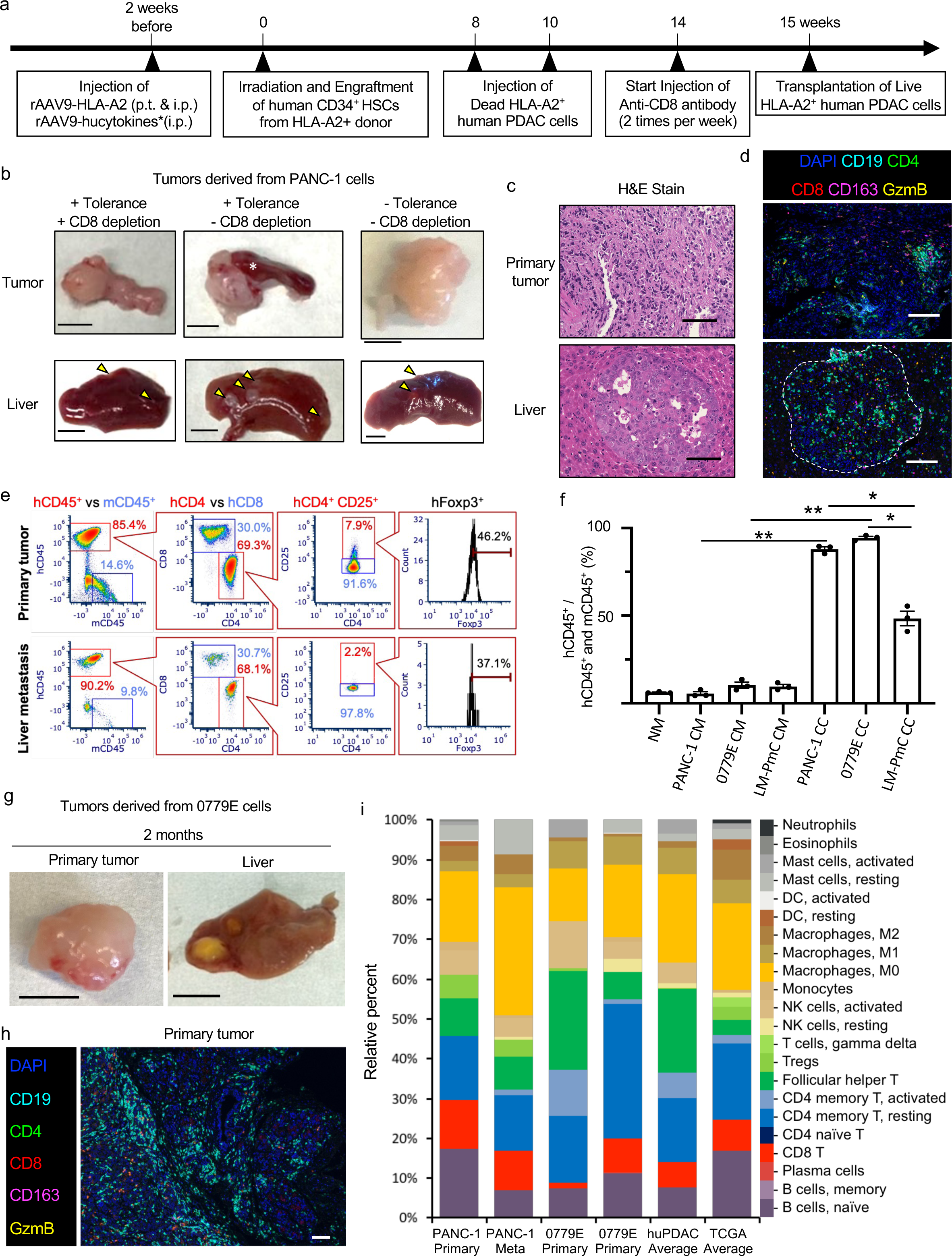
Establishment of patient derived huPDAC mice. (a) A scheme showing the timeline of huPDAC mouse development. Genes encoding HLA and huCytokines are introduced into NSG mice that lack mouse β2-microglobulin by AAV9-mediated gene transfer. Mice are irradiated and engrafted with cord-derived HSCs. To induce B cell tolerance, UV-irradiated PDAC cells were injected 8 and 10 weeks after HSC transplantation. To deplete CD8^+^ T cells, an anti-CD8 Ab was injected twice per week for 2 weeks from 14 weeks after HSC transplantation. PDAC cells were orthotopically implanted into mice 15 week-post HSC transplantation. (b) Macroscopic appearance of the PDAC and liver of PANC-1 huPDAC mice with or without B cell tolerance and CD8 depletion. Arrowheads, metastases; *, spleen. Scale bar, 5 mm. (c) Representative H&E images of PDAC and liver sections prepared from huPDAC mice. Scale bar, 100 μm. (d) Representative multiplex immunofluorescence images of PDAC and liver sections prepared from PANC-1 huPDAC mice. Scale bar, 100 μm. Blue, DAPI; cyan, CD19; green, CD4; red, CD8; magenta, CD163; yellow, GZmB. (e) Flow cytometry showing human versus mouse CD45^+^ cells, and human CD4, CD8, and CD4^+^ CD25^+^ Foxp3^+^ (Tregs) cells isolated from primary and metastatic PANC-1 huPDAC tumors. (f) *In vitro* immune cell cultures. Splenocytes harvested from HIS mice were cultured in normal media (NM) or conditioned media (CM) prepared from the indicated PDAC cells, or co-cultured with the PDAC cells (CC). *p<0.05, **, p<0.01. (g) Primary and metastatic 0799E huPDAC tumors 1 and 2 months after implantation. Scale bar, 5 mm. (h) A multiplex immunofluorescence image of a primary 0799E huPDAC tumor. Scale bar, 100 μm. (i) CIBERSORTx analysis showing the proportion of each immune cell fraction in huPDAC tumors in comparison with cell fractions in TCGA patient-derived PDAC samples. More than 65% of PBMCs were reconstituted by hCD45^+^ cells in the HIS mice.

To grow an HLA-matched PDAC in our HLA-A2^+^ HIS mice, we first searched for HLA-A2^+^ human PDAC cells. Consistent with earlier reports^20^, HLA genotyping showed that *Kras^G12D^* PANC-1 human PDAC cells expressed HLA-A2 (Supplementary Fig. 1a). Additional HLAs, such as A11, B38, and DRB1*13, were also detected. The expression of HLA-A2 protein was verified by flow cytometry (Supplementary Fig. 1b). Basic *in vivo* tumorigenicity of the PANC-1 cells was confirmed by orthotopic implantation of the cells into nude mice

(Supplementary Fig. 2). Upon growing PANC-1 tumors in our HIS mice, we had a concern that the tumors would be rejected given that the cells expressed additional HLAs other than HLA-A2. We therefore employed several strategies to facilitate engraftment of the PANC-1 cells by inducing tolerance to B cells that mediate humoral immunity and by depleting CD8^+^ cytotoxic T cells during the tumorigenic phase (Fig. 1a). To induce B cell tolerance, apoptotic PANC-1 cells were injected subcutaneously into the flank of HIS mice 8-10 weeks after HSC transplantation, right around the timing when B cell reconstitution occurred^12^. CD8^+^ T cells were depleted by intraperitoneal injections of an anti-CD8 antibody (Ab) starting 14 weeks after HSC transplantation and 1 week before T cell maturation^12^ and orthotopic implantation of PANC-1 cells. Despite our concern, PANC-1 cells consistently formed ∼10 mm orthotopic tumors within 8 weeks in all the HIS mice, including those that did not receive any treatment to induce B cell tolerance or CD8^+^ T cell depletion (Fig. 1b, upper panels). All the huPDAC mice developed multiple liver metastases (Fig. 1b, lower panels). Histology revealed fibrotic stroma in the primary tumor and multiple micrometastases in the liver (Fig. 1c).

Multiplex immunofluorescence showed that both the primary tumors and liver metastases were infiltrated by a variety of human immune cells, such as CD19^+^ B cells, CD4^+^ T cells, CD8^+^ T cells, CD163^+^ monocytes, and granzyme B^+^ cytotoxic T and natural killer (NK) cells (Fig. 1d). These results show that an established HLA-A2^+^ human PDAC cell line can be grown in our HIS mice, and that various human immune cells infiltrate both the resulting primary and metastatic tumors.

Flow cytometry of tumor-infiltrating lymphocytes (TILs) confirmed the presence of hCD45^+^ immune cells in both the primary and metastatic tumors in the liver (Fig. 1e). Approximately 90% of the TILs were of human origin, in line with the high proportion of hCD45^+^ cells in the PBMCs at the time of tumor implantation (refer to mouse #2, Supplementary Table 1).

Within the hCD45^+^ cells, CD4^+^ and CD8^+^ T cells existed at a 7:3 ratio similar to patient-derived PDAC samples^21^. CD4^+^ CD25^+^ Foxp3^+^ human regulatory T cells (Tregs)^22^ were also detected in the tumors. Interestingly, the CD45^+^ cells in the tumors were mostly of human origin even when only <35% of the PBMCs were reconstituted by hCD45^+^ cells in the original HIS mice (mouse #3-5, Supplementary Table 1). In contrast, the CD45^+^ cells in non-tumor tissues such as the spleen and peripheral circulation were mainly of mouse origin.

These results suggest that the PDAC microenvironment contributes to the maintenance of TILs. To test this possibility, we cultured splenocytes harvested from HIS mice in the presence or absence of PDAC cells. Approximately 70% of the CD45^+^ cells were of human origin, and the remaining were mouse origin at the time they were harvested from the mice (Supplementary Fig. 3). When the cells were cultured in normal media, the proportion of hCD45^+^ cells decreased to 10% allowing mouse CD45^+^ cells to dominate (Fig. 1f, Supplementary Fig. 4). On the other hand, >60% of the CD45^+^ cells were of human origin when the cells were co-cultured with PANC-1 cells. Co-culturing the CD45^+^ cells with 0779E HLA-A2^+^ patient-derived primary human PDAC cells^23, 24^ (Supplementary Fig. 5) led to an even higher proportion of hCD45^+^ cells. Direct contact between the CD45^+^ cells and human PDAC cells appeared important because conditioned medium prepared from the PDAC cells had only a minor effect on the maintenance of hCD45^+^ cells. Of note, LM-PmC mouse PDAC cells derived from transgenic *Kras-LSL^GD^*^12^*, p53-LSL^172H^, Pdx-1-cre* mice with a C57BL/6J and 129S1/SvImJ hybrid background^25^ had a weak effect on the maintenance of hCD45^+^ cells. These results suggest that species-specific interaction between TILs and PDAC cells drives the formation of the PDAC TIME.

We next studied whether 0779E, a primary PDAC cell line, could be grown in our HIS mice. 0799E cells grew in nude mice confirming its tumorigenicity (Supplementary Fig. 6). Similar to PANC-1 cells, 0799E cells formed orthotopic tumors in our HIS mice within 4 weeks without the need for B cell tolerance or depletion of CD8^+^ T cells (Supplementary Fig. 7).

Liver metastasis was not observed under this condition. However, liver metastases were observed when 10 times fewer 0779E cells were injected to allow the primary tumors to slowly grow for up to 2 months (Fig. 1g). Multiplex immunofluorescence revealed the infiltration of various human immune cells in PDAC tissue (Fig. 1h). These results indicate that patient-derived xenograft (PDX) PDAC models can be generated in our HIS mice.

To further analyze how well the huPDAC tumors recapitulate the TIME of patient-derived PDAC samples, we performed a bulk RNA-seq analysis of the huPDAC tumors which were highly reconstituted with human CD45^+^ cells in reference to 182 patient-derived PDAC samples available in The Cancer Genome Atlas (TCGA) database (https://www.cancer.gov/tcga). Analysis of cell fractions, inferred from RNA-seq, using CIBERSORTx^26, 27^, showed that the immune cell fractions in the huPDAC tumors and the TCGA patient samples were similar (Fig. 1i, Supplementary Table 2). Cluster analysis showed that both PANC-1 and 0779E tumors clustered with the patient samples in the TCGA database (Supplementary Fig. 8). The gapstat statistic showed that the optimal number of clusters is one, indication similarity between the groups. Multidimensional Scaling also found the huPDAC tumors to be similar to TCGA (data not shown). Wilcoxon rank analysis showed the cell fractions in huPDAC tumors and TCGA to be similar in general, except that some small differences were noted in immune cells, such as NK cells, T follicular helper cells, CD4^+^ memory resting cells, and M2 macrophages, which can contribute to humoral immunity.

As a proof-of-principle experiment to assess the utility of the huPDAC mice, we studied the effect of the iRGD peptide on their TIME. iRGD (amino acid sequence: CRGDKGPDC) is a tumor-penetrating peptide that delivers drugs deep into the extravascular tumor tissue by targeting αvβ3/β5 integrins and neuropilin-1 (NRP-1)^28, 29^. iRGD induces a NRP-1-dependent transcellular transport mechanism in the tumor that propels the peptide and co-injected molecules through the vascular barrier and subsequent tumor cell layers^30^. This mechanism allows iRGD to deliver co-injected drugs into tumors, thereby providing a versatile drug delivery platform for cancer therapy^31^. iRGD has shown a remarkable preliminary efficacy as a tumor-specific enhancer of gemcitabine + nab-paclitaxel therapy in stage 4 PDAC patients in a phase 1b trial^32^, and is now in multiple phase 2 studies (e.g., ASCEND study; NCT05042128). We have recently found that iRGD not only delivers drugs, but also modifies the TIME in syngeneic PDAC mice^33^. Long-term treatment with iRGD increases the ratio of CD8^+^ T cells over Tregs in PDAC tissue, and sensitizes PDAC to an ICB. iRGD targets Tregs selectively in PDAC tissue because these tumor-resident Tregs, but not those in other tissues, express the αvβ5 integrin in addition to NRP-1. Both PANC-1 and 0799E huPDAC mice harbor αvβ5 integrin^+^ and NRP-1^+^ human Tregs in the tumors but not in the spleen (Supplementary Fig. 9) providing an opportunity to test the immunomodulatory effects of iRGD in a relevant preclinical setting.

Systemic treatment of PANC-1 huPDAC mice with iRGD led to an increase in the CD8/Treg ratio in the PDAC tissue based on multiplex immunofluorescence (Fig. 2a and b) and flow cytometry (Supplementary Fig. 10). The treatment did not affect ratios in the spleen consistent with the tumor-specific effect of iRGD (Fig. 2c). CIBERSORTx was used to infer immune fractions of both iRGD-treated and untreated samples from RNA-seq data (Fig. 2d). Hierarchical clustering of immune cell fractions revealed that iRGD-treated huPDAC tumors were clustered separately from untreated huPDAC tumors, suggesting that iRGD treatment altered the TIME (Fig. 2e). The CD8/Treg ratio numerically increased upon iRGD treatment based on CIBERSORTx analysis, but the difference did not reach statistical significance with the limited number of animals tested (Figs. 2d, 2f, and Supplementary Table. 3).

**Figure 2.**
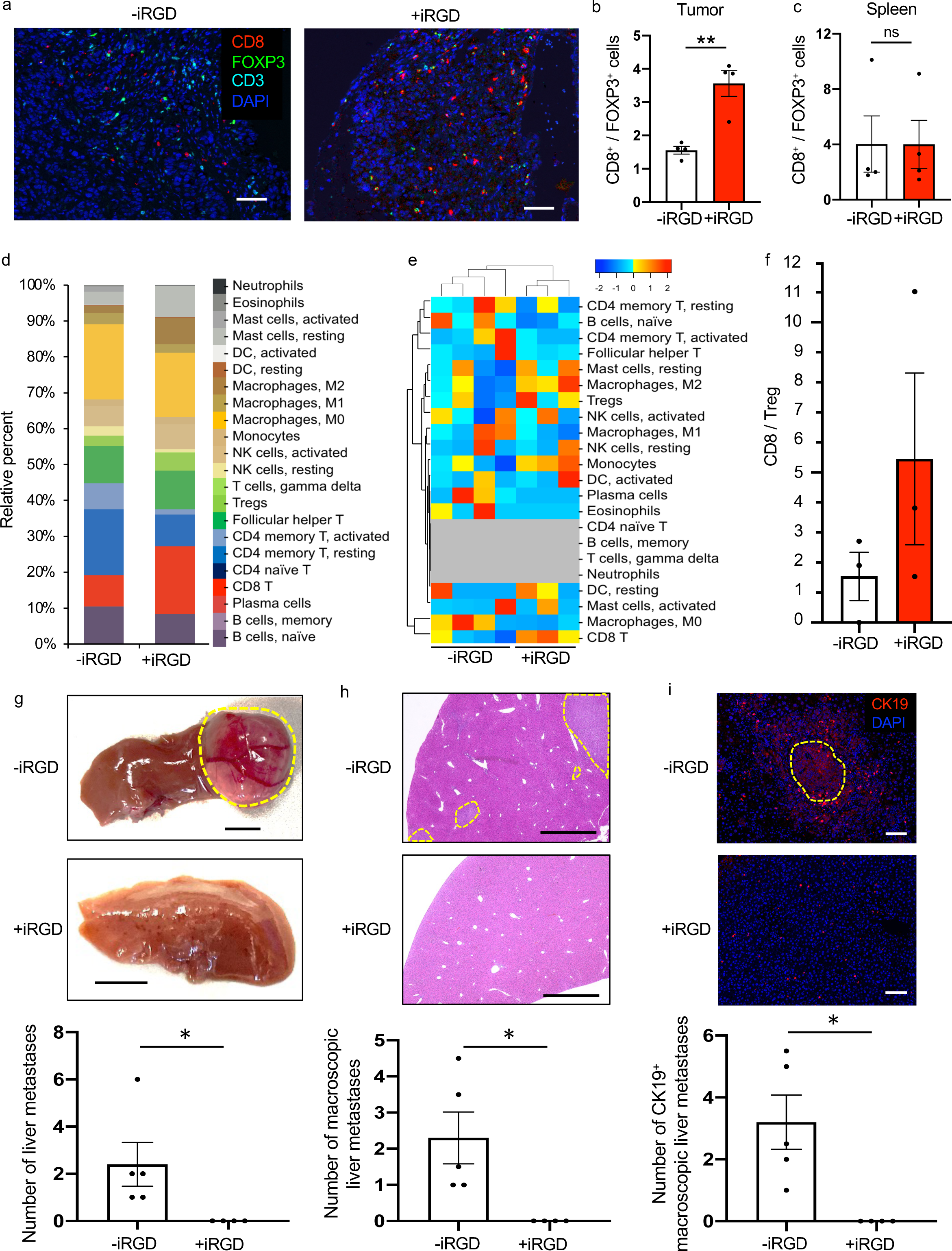
The effect of systemic iRGD treatment on huPDAC tumors. huPDAC mice bearing an orthotopic PANC-1 tumor were treated with or without 1.2 mg / kg of iRGD 3 times a week for 8 weeks. (a-c) The tumors and spleen were subjected to immunofluorescence at the end of the treatment study. Representative immunofluorescence images of PDAC sections are shown in (a). Red, CD8; green FOXP3; blue, DAPI. Scale bar, 100 μm. The ratio of CD8^+^ T cells over FoxP3^+^ Tregs in the PDAC (b) and spleen (c) was quantified by counting the cells in 5 random high-power fields per section for 4 mice per arm. Error bars, mean ± standard error: **p < 0.01. (d-f) The PDAC tissue was also subjected to RNA-seq. Hierarchical clustering of major immune cells in the PDAC tissue is shown in (d). Samples of untreated 0779E huPDAC tumors were also included in the analysis. CIBERSORTx analysis showing the proportion of each immune cell fraction in huPDAC tumors is shown in (e). The CD8/Treg ratio in the huPDAC tumors was also analyzed by CIBERSORTx (f). (g-i) The effect of systemic iRGD monotherapy on liver metastasis in PDAC-1 huPDAC mice was characterized based on the macroscopic appearance of the liver (g) and microscopic analyses of the liver sections stained with H&E (h) or an anti-CK19 antibody (red) (i). Dotted circles: metastases. Blue (i), DAPI. Scale bars; 5 mm (g), 500 μm (h), and 100 μm (i). The number of liver metastases under each analytical condition was counted either macroscopically or under a microscope as summarized in the bar diagrams. Error bars, mean ± standard error: *p < 0.05. Scale bar, 100 μm.

In line with historical data^31, 34, 35^, iRGD monotherapy did not inhibit the growth of primary tumors (Supplementary Fig. 11). However strikingly, iRGD significantly inhibited liver metastasis in the PANC-1 huPDAC mice. There were no visible nodules on the liver surface (Fig. 2g, Supplementary Fig. 12) or microscopically detectable tumors in serial liver sections stained with hematoxylin and eosin (H&E) (Fig. 2h, Supplementary Fig. 13) and no micrometastases were observed after CK19 immunostaining (Fig. 2i). The effect of iRGD on metastasis is consistent with our previous observation made in traditional tumor mouse models produced with immunodeficient mice^36^. However, the effects are significantly more pronounced in the huPDAC mice implicating a role of immunity in the anti-metastatic effects of iRGD.

This report introduces a novel humanized PDAC mouse model, which develops an orthotopic human PDAC that spontaneously metastasizes to the liver. The tumors are infiltrated by a full panel of human immune cells that are found in patient-derived PDAC samples recapitulating the TIME of human PDAC tissue. Patient-derived primary PDAC cells as well as established human PDAC cell lines can be used to develop the huPDAC mice, providing a versatile model system to study the origin of the PDAC TIME and the response of the TIME to immunomodulators, such as iRGD, to enhance development of novel immunotherapies and personalized cancer therapy.

## Methods

### Tumor cells

PANC-1 cells were purchased from AATC (Manassas, VA). 0799E cells were established from primary human PDAC tissue. LM-PmC (mCherry-labeled LM-P PDAC cells) was established from liver metastases in Pdx1-Cre; KrasLSL-G12D/+; Trp53LSL-R172H/+ mice on a B6/129 background^34^. PANC-1 and 0799E cells were authenticated using eighteen mouse short tandem repeat performed by ATCC. HLA typing of PANC-1 and 0779E cells was performed by Histogenetics LLC (Ossining, NY).

### Mice

All animal experiments were performed according to procedures approved by the Institutional Animal Care and Use Committee (IACUC) at Columbia University (New York, NY). Athymic nude mice were purchased from Envigo (Indianapolis, IN). NOD.Cg-*B2m^tm1Unc^ Prkdc^scid^ Il2rg^tm1Wjl^*/SzJ (NSG-B2m) triple mutant mice with combined features of severe combined immune deficiency mutation, IL-2 receptor γ chain deficiency, and β2-microglobulin (β2m) deficiency, were purchased from The Jackson Laboratories (Bar Harbor, ME). The mice were maintained under specific pathogen-free conditions in the animal facilities at the Columbia University Medical Center.

### Generation of HIS mice

Three-week old NSG-B2m mice were given an intravenous (i.v.) injection of recombinant adeno-associated virus serotype 9 AAV9 encoding for human IL-3, IL-15, and GM-CSF, as well as peri-thoracic injection of AAV9 encoding for HLA-A2. One to two weeks after the transduction, the mice were exposed to 1.5 Gy of whole-body sublethal irradiation for myeloablation. Four hours later, each mouse received an i.v. injection of 1 × 10^5^ HLA-A2^+^ matched CD34^+^ human HSCs, which were isolated from human fetal liver, as previously described^12, 37^.

### Tumor mouse models

PDAC in nude mice were produced by orthotopic pancreatic injections of 1 x 10^6^ PANC-1 or 0779E cells under deep anesthesia. huPDAC mice were generated by orthotopic pancreatic injections of 1 x 10^7^ PANC-1 or 1 x 10^6^-10^7^ 0779E cells into HIS mice under deep anesthesia. In some cases, HIS mice were subcutaneously injected with apoptotic PANC-1 cells which were exposed UV 60 min 8-10 weeks after HSC transplantation to induce B cell tolerance before orthotopic PANC-1 implantation. CD8^+^ T cells were transiently depleted in some HIS mice prior to orthotopic PANC-1 implantation by intraperitoneally injecting 4 mg/kg anti-CD8 Ab (NHP Reagent Resource, #MT807R1; Worcester, MA) 2 times a week for 3 weeks starting 14 weeks after HSC transplantation. To check the expression of HLA-A2, PDAC cells were resuspended in 0.1% bovine serum albumin in PBS, and stained with the following Abs for flow cytometry: HLA-A2 (BioLegend #343306; San Diego, CA), Zombie NIR™ Fixable Viability Kit (BioLegend #423106).

### HLA Typing

HLA typing was performed by Histogenetics LLC (Ossining, NY). Sequencing was performed on the Illumina platform. Allele database version is 3.39.0 (Jan 2020).

### Mouse treatment studies

Systemic treatment studies in huPDAC mice were performed as follows. Eight days after orthotopic tumor cell inoculation, the mice were randomized into two treatment arms, phosphate buffered saline (PBS) and iRGD (1.2 mg / kg). The iRGD peptide (cyclic acetyl-CRGDKGPDC-NH2) was purchased from a commercial vendor (LifeTein, Hillsborough, NJ). The treatment was given 3 times a week for 8 weeks. The tumors and major organs were harvested at the end of the studies, weighed, and processed for flow cytometry, multiplex immunofluorescence, H&E staining, and RNA-seq. The treatment studies were terminated according to the guidelines by the IACUC at Columbia University.

### Isolation of immune cells from PDAC tissue

Tumor-infiltrating immune cells were isolated as previously described^33^. In brief, 2-4 mm pieces of tumors from huPDAC mice were treated with an enzyme mix from a tumor dissociation kit (Miltenyi Biotec, Bergisch Gladbach, Germany). The tissues were dissociated using a gentle MACS Dissociator (Miltenyi Biotec) and incubated for 40 min at 37 °C. Debris was then removed using a debris removal solution (Miltenyi Biotec). Splenocytes were prepared by gently grinding the spleen, passing the cell suspension through a 40 μm cell strainer, and lysing the red cells with an ACK lysing buffer (Thermo Fisher Scientific, Waltham, MA). The isolated cells were resuspended in 0.1% bovine serum albumin in PBS, and stained with the following Abs for flow cytometry: human CD4 (BioLegend, #300532), human CD3 (BioLegend, #317324), mouse CD45 (BioLegend #103130), human CD8 (BioLegend #344712), human CD45 (BioLegend #304014), human CD25 (BioLegend #302606), human FOXP3 (BioLegend #320116), Alexa Fluor® 647-conjugated anti-mouse integrin αvβ5 Ab (clone ALULA; BD Biosciences), Zombie NIR™ Fixable Viability Kit (BioLegend #423106) and mouse BD Fc Block™ (BD Biosciences #553142; San Jose, CA).

### Histological Analysis

Tumor and liver tissue samples were fixed in 4% PFA/PBS and embedded in paraffin. Using 4 μm sections of the samples, the tissues were subjected to H&E as previously performed^34^. Images were acquired by Keyence BZ-X800 (Keyence, Itasca, IL). The sections were also subjected to multiplex immunofluorescence, which was performed using an Opal 6-Plex Detection Kit (Akoya Biosciences, Menlo Park, CA) with the following Abs: Anti-human Abs against CD19 (Leica #NCL-L-CD19-163; Biosystems, Buffalo Grove, IL), CD8 (Leica #PA0183), CD163 (Leica #PA0090), CD4 (Abcam #ab133616), GzmB (Leica #PA0291). CD8 (4B11) (Leica #PA0183 and #CD8-4B11-L-CE), CD4 (EPR6855) (Abcam #ab133616) and FoxP3 (236A/E7) (Abcam #ab20034), CD3 (LN10) (Leica #CD3-565-L-CE), and CK19 (Abcam #ab53119). The sections were also stained with DAPI. Images were acquired by Akoya’s vectra polaris automated quantitative pathology imaging system (Akoya Bioscience). For conventional immunofluorescence, antigen retrieval techniques were applied human tissue sections using IHC Antigen Retrieval Solution (00-4956-58, Invitrogen). These tissues were stained with Abs against human CD4 pre-labeled with Alexa Fluor 488 using an Ab labeling kit (Molecular Probes, A20187), human/mouse Foxp3 (clone 1054c; Novus Biologicals, Centennial, CO) pre-labeled with Alexa Fluor 555 using an Ab labeling kit (Molecular Probes, A20181), Alexa Fluor® 647-conjugated anti-mouse integrin αvβ5 Ab (clone ALULA; BD Biosciences), human NRP-1 (AD5-17F6; Milteny Biotec) pre-labeled with Alexa Fluor 647 using an Ab labeling kit (Molecular Probes, A20186). Images were acquired with an LSM 710 confocal microscope (Carl Zeiss, Oberkochen, Germany). A ZEN 3.0 SR black edition software (Zeiss) was used for acquisition and analysis of the images.

### *In vitro* immune cell survival assay

Splenocytes isolated from the spleen of HIS mice were cultured on a monolayer of PANC-1, 0799E or LM-PmC cells prepared 3 days before, or cultured either in normal DMEM containing 10% FBS or in conditioned media prepared from 3-day-old cultures of each PDAC cell type. Three days later, the immune cells were harvested, and survival was analyzed by flow cytometry using the following antibodies. Mouse CD45 (BioLegend #103130), human CD45 (BioLegend #304014), Zombie NIR™ Fixable Viability Kit (BioLegend #423106) and mouse BD Fc Block™ (BD Biosciences #553142; San Jose, CA). Live single cell populations were determined using forward and side scatters combined with Zombie NIR. The proportion of human and mouse CD45^+^ cells was studied.

### RNA-seq

Tumors were snap frozen in liquid nitrogen and stored at -80°C. Total RNA was extracted from the samples using miRNeasy kits (Qiagen; Hilden, Germany). RNA integrity was assessed with an Agilent Bioanalyzer using an RNA Pico kit (Agilent Technologies, Palo Alto, CA). For STRDPOLYA, a poly-A pull-down was used to enrich mRNAs from the total RNA samples, and cDNA libraries were constructed using Illumina TruSeq chemistry. The libraries were fragmentated, ligated to an adapter, and sequenced using an Illumina NovaSeq 6000 (Illumina Inc, San Diego, CA) at the Columbia Genome Center. Multiplex samples were used in each lane to yield targeted number of paired-end 100 bp reads for each sample.

Numbers of samples: PANC-1 primary, iRGD treated, n=4; PANC-1 primary, iRGD untreated, n=3; 0799E primary, iRGD untreated, n=2; PANC=1, liver metastasis, iRGD untreated, n=1. Data was deposited in the Gene Expression Omnibus (GEO)^38^, GSE222189.

### Bioinformatic analysis

Relative proportions of each immune cell population in huPDAC tumors and patient-derived human PDAC samples from TCGA database (TCGAPAN) were inferred from the RNA-seq data using CIBERSORTx^26, 27^ The immune cell composition in untreated huPDAC tumors and TCGA human PDAC samples, and the composition in untreated versus iRGD-treated huPDAC tumors were compared. The LM22 immune cell signature matrix (with updated gene symbols) was used^39^. Significance analysis was based on 1000 permutations. The B batch adjustment method was used to correct for differences in the method of the current experimental data (RNASeq) and the method used to derive the signature matrix (microarray). Each group of cells that was compared to other groups of cells was submitted to CIBERSORTx separately. Transcripts per million (TPM) values for HIS samples were obtained from FASTQ files using Kallisto^40^ with Ensembl v96 annotation, based upon the GRCm38.p6 assembly of the mouse genome. TPM values for TCGA files were obtained by converting TCGA count files. All subsequent analyses on the cell fraction data were performed in R^41^. Hierarchical clustering of cell fractions was performed and dendrograms and heatmaps were generated with the heatmap.2 command in gplots^42^. A Euclidean distance metric and average linkage clustering was used^43^. The optimal number of clusters was determined with the gapstat statistic^44^ as implemented in NbClust^45^. Multidimensional scaling^46^ was performed using the cmdscale command in the R stats package^41^. Differences between conditions on a cell type level were analyzed by the Wilcoxon rank test^47^ and the Benjamini-Hochberg false discovery rate^48^.

### Statistical analyses

Statistical analysis was performed using two-tailed unpaired student’s t-test between two groups and multiple comparison test after Ryan’s method in over three groups, except as noted above.

### Author information

Norio Miyamura, Kodai Suzuki, Yuki Kunisada and Kazuki N Sugahara

**Biomedical Informatics Shared Resource, Herbert Irving Comprehensive Cancer Center, Columbia University Irving Medical Center, New York, NY USA.**

Richard Friedman and Aristidis Floratos

**Department of Biomedical Informatics, Columbia University Irving Medical Center, New York, NY USA.**

Richard Friedman and Aristidis Floratos

**Department of Systems Biology, Columbia University Irving Medical Center, New York, NY USA.**

Aristidis Floratos

**Aaron Diamond AIDS Research Center, Division of Infectious Diseases, Department of Medicine, Vagelos College of Physicians and Surgeons, Columbia University Irving Medical Center, New York, NY USA.**

Kazuya Masuda and Moriya Tsuji

**Department of Surgery, Division of Surgical Oncology, Moores Cancer Center, University of California, San Diego, La Jolla, CA, USA.**

Andrew M. Lowy

## Acknowledgements

We thank Yoko Odagiri for assisting the experiments. We thank Aaron Newman for helpful correspondence. We also thank the Columbia Genome Center, Biomedical Informatics, Flow Cytometry Microscopy and Shared Equipment Core, Molecular Pathology/MPSR Core, and the Human Immune Monitoring Core at Columbia University. This work was supported by grants R01CA167174 (K.N.S.) and R01CA155620 (A.M.L.) from the National Cancer Institute of NIH, R21AI151469 (M.T.) from the National Institute of Allergy and Infectious Diseases of NIH, the Translational Research Grant from the Pancreatic Cancer Action Network (K.N.S.), Idea Award with Special Focus from the Department of Defense (K.N.S.), the Alexandrina M. McAfee Trust Foundation (A.M.L) and the Research for a Cure of Pancreatic Cancer Fund (A.M.L.). K.S. was supported by the Uehara Memorial Foundation Research Fellowship (#201941078, Japan). The results shown here are in whole or part based upon data generated by the TCGA Research Network: https://www.cancer.gov/tcga.

## Contributions

N.M., M.T. and K.N.S. developed the concept of the study. N.M. performed most of the experiments. K.S. and Y.K. assisted with flow cytometry. R.F. and A.F. performed the RNA-seq analysis. K.M. prepared the HIS mice. A.M.L. developed the 0799E cells. N.M. and K.N.S. wrote the manuscript. M.T. and K.N.S. supervised the study.

## Reference

1 Wei, S. C., Duffy, C. R. & Allison, J. P. Fundamental Mechanisms of Immune Checkpoint Blockade Therapy. Cancer Discovery 8, 1069–1086, doi:10.1158/2159-8290.Cd-18-0367 (2018).

2 Pointer, K. B., Pitroda, S. P. & Weichselbaum, R. R. Radiotherapy and immunotherapy: open questions and future strategies. Trends in Cancer 8, 9–20, doi:https://doi.org/10.1016/j.trecan.2021.10.003 (2022).

3 Chow, A., Perica, K., Klebanoff, C. A. & Wolchok, J. D. Clinical implications of T cell exhaustion for cancer immunotherapy. Nature Reviews Clinical Oncology 19, 775–790, doi:10.1038/s41571-022-00689-z (2022).

4 Sharma, P., Hu-Lieskovan, S., Wargo, J. A. & Ribas, A. Primary, Adaptive, and Acquired Resistance to Cancer Immunotherapy. Cell 168, 707–723, doi:https://doi.org/10.1016/j.cell.2017.01.017 (2017).

5 Sugiyama, D., et al. Anti-CCR4 mAb selectively depletes effector-type FoxP3+CD4+ regulatory T cells, evoking antitumor immune responses in humans. Proceedings of the National Academy of Sciences 110, 17945–17950, doi:doi:10.1073/pnas.1316796110 (2013).

6 Kubli, S. P., Berger, T., Araujo, D. V., Siu, L. L. & Mak, T. W. Beyond immune checkpoint blockade: emerging immunological strategies. Nature Reviews Drug Discovery 20, 899–919, doi:10.1038/s41573-021-00155-y (2021).

7 Rios-Doria, J., Stevens, C., Maddage, C., Lasky, K. & Koblish, H. K. Characterization of human cancer xenografts in humanized mice. Journal for ImmunoTherapy of Cancer 8, e000416, doi:10.1136/jitc-2019-000416 (2020).

8 Allen, T. M., et al. Humanized immune system mouse models: progress, challenges and opportunities. Nature Immunology 20, 770–774, doi:10.1038/s41590-019-0416-z (2019).

9 Horowitz, N. B., et al. Humanized Mouse Models for the Advancement of Innate Lymphoid Cell-Based Cancer Immunotherapies. Frontiers in Immunology 12, doi:10.3389/fimmu.2021.648580 (2021).

10 Tian, H., Lyu, Y., Yang, Y.-G. & Hu, Z. Humanized Rodent Models for Cancer Research. Frontiers in Oncology 10, doi:10.3389/fonc.2020.01696 (2020).

11 Tsuji, M. & Akkina, R. Editorial: Development of Humanized Mouse Models for Infectious Diseases and Cancer. Frontiers in Immunology 10, doi:10.3389/fimmu.2019.03051 (2020).

12 Huang, J., Li, X., Coelho-dos-Reis, J. G. A., Wilson, J. M. & Tsuji, M. An AAV Vector-Mediated Gene Delivery Approach Facilitates Reconstitution of Functional Human CD8+ T Cells in Mice. PLOS ONE 9, e88205, doi:10.1371/journal.pone.0088205 (2014).

13 Huang, J., et al. Human immune system mice immunized with Plasmodium falciparum circumsporozoite protein induce protective human humoral immunity against malaria. Journal of Immunological Methods 427, 42–50, doi:https://doi.org/10.1016/j.jim.2015.09.005 (2015).

14 Huang, J., et al. Targeted Co-delivery of Tumor Antigen and α-Galactosylceramide to CD141+ Dendritic Cells Induces a Potent Tumor Antigen-Specific Human CD8+ T Cell Response in Human Immune System Mice. Frontiers in Immunology 11, doi:10.3389/fimmu.2020.02043 (2020).

15 Li, X., et al. A potent adjuvant effect of a CD1d-binding NKT cell ligand in human immune system mice. Expert Review of Vaccines 16, 73–80, doi:10.1080/14760584.2017.1256208 (2017).

16 Li, X., et al. Human CD8+ T cells mediate protective immunity induced by a human malaria vaccine in human immune system mice. Vaccine 34, 4501–4506, doi:https://doi.org/10.1016/j.vaccine.2016.08.006 (2016).

17 Nogueira, R. T., Sahi, V., Huang, J. & Tsuji, M. Human IgG repertoire of malaria antigen-immunized human immune system (HIS) mice. Immunology Letters 188, 46–52, doi:https://doi.org/10.1016/j.imlet.2017.06.001 (2017).

18 Sharma, A., et al. Respiratory Syncytial Virus (RSV) Pulmonary Infection in Humanized Mice Induces Human Anti-RSV Immune Responses and Pathology. Journal of Virology 90, 5068–5074, doi:doi:10.1128/JVI.00259-16 (2016).

19 Coelho-Dos-Reis, J. G. A., et al. Functional Human CD141+ Dendritic Cells in Human Immune System Mice. The Journal of Infectious Diseases 221, 201–213, doi:10.1093/infdis/jiz432 (2019).

20 Zhu, K. et al. HLA-A0201 positive pancreatic cell lines: new findings and discrepancies. Cancer Immunology, Immunotherapy 56, 719–724, doi:10.1007/s00262-006-0217-8 (2007).

21 Tímár, J., et al. Neoadjuvant Immunotherapy of Oral Squamous Cell Carcinoma Modulates Intratumoral CD4/CD8 Ratio and Tumor Microenvironment: A Multicenter Phase II Clinical Trial. Journal of Clinical Oncology 23, 3421–3432, doi:10.1200/JCO.2005.06.005 (2005).

22 Dikiy, S. & Rudensky, A. Y. Principles of regulatory T cell function. Immunity 56, 240–255, doi:https://doi.org/10.1016/j.immuni.2023.01.004 (2023).

23 Fujimura, K., et al. A hypusine-eIF5A-PEAK1 switch regulates the pathogenesis of pancreatic cancer. Cancer Res 74, 6671–6681, doi:10.1158/0008-5472.Can-14-1031 (2014).

24 Scully, K. M., et al. E47 Governs the MYC-CDKN1B/p27(KIP1)-RB Network to Growth Arrest PDA Cells Independent of CDKN2A/p16(INK4A) and Wild-Type p53. Cell Mol Gastroenterol Hepatol 6, 181–198, doi:10.1016/j.jcmgh.2018.05.002 (2018).

25 Tseng, W. W., et al. Development of an Orthotopic Model of Invasive Pancreatic Cancer in an Immunocompetent Murine Host. Clinical Cancer Research 16, 3684–3695, doi:10.1158/1078-0432.Ccr-09-2384 (2010).

26 Newman, A. M., et al. Robust enumeration of cell subsets from tissue expression profiles. Nature methods 12, 453–457, doi:10.1038/nmeth.3337 (2015).

27 Newman, A. M., et al. Determining cell type abundance and expression from bulk tissues with digital cytometry. Nature biotechnology 37, 773–782, doi:10.1038/s41587-019-0114-2 (2019).

28 Sugahara, K. N., et al. Tissue-Penetrating Delivery of Compounds and Nanoparticles into Tumors. Cancer Cell 16, 510–520, doi:https://doi.org/10.1016/j.ccr.2009.10.013 (2009).

29 Teesalu, T., Sugahara, K. N., Kotamraju, V. R. & Ruoslahti, E. C-end rule peptides mediate neuropilin-1-dependent cell, vascular, and tissue penetration. Proceedings of the National Academy of Sciences 106, 16157–16162, doi:10.1073/pnas.0908201106 (2009).

30 Pang, H.-B., et al. An endocytosis pathway initiated through neuropilin-1 and regulated by nutrient availability. Nature Communications 5, 4904, doi:10.1038/ncomms5904 (2014).

31 Sugahara, K. N., et al. Coadministration of a Tumor-Penetrating Peptide Enhances the Efficacy of Cancer Drugs. Science 328, 1031–1035, doi:10.1126/science.1183057 (2010).

32 Dean, A., et al. Dual αV-integrin and neuropilin-1 targeting peptide CEND-1 plus nab-paclitaxel and gemcitabine for the treatment of metastatic pancreatic ductal adenocarcinoma: a first-in-human, open-label, multicentre, phase 1 study. The Lancet Gastroenterology & Hepatology 7, 943–951, doi:https://doi.org/10.1016/S2468-1253(22)00167-4 (2022).

33 Suzuki, K. et al. Tumor-resident regulatory T cells in pancreatic cancer express the αvβ5 integrin as a targetable activation marker. (submitted).

34 Hurtado de Mendoza, T., et al. Tumor-penetrating therapy for β5 integrin-rich pancreas cancer. Nature Communications 12, 1541, doi:10.1038/s41467-021-21858-1 (2021).

35 Akashi, Y., et al. Anticancer effects of gemcitabine are enhanced by co-administered iRGD peptide in murine pancreatic cancer models that overexpressed neuropilin-1. British Journal of Cancer 110, 1481–1487, doi:10.1038/bjc.2014.49 (2014).

36 Sugahara, K. N., et al. Tumor-Penetrating iRGD Peptide Inhibits Metastasis. Molecular Cancer Therapeutics 14, 120–128, doi:10.1158/1535-7163.MCT-14-0366 (2015).

37 Nauman, G., et al. Defects in Long-Term APC Repopulation Ability of Adult Human Bone Marrow Hematopoietic Stem Cells (HSCs) Compared with Fetal Liver HSCs. The Journal of Immunology 208, 1652–1663, doi:10.4049/jimmunol.2100966 (2022).

38 Edgar, R., Domrachev, M. & Lash, A. E. Gene Expression Omnibus: NCBI gene expression and hybridization array data repository. Nucleic Acids Res 30, 207–210, doi:10.1093/nar/30.1.207 (2002).

39 Newman, A. M., et al. Robust enumeration of cell subsets from tissue expression profiles. Nature Methods 12, 453–457, doi:10.1038/nmeth.3337 (2015).

40 Bray, N. L., Pimentel, H., Melsted, P. & Pachter, L. Near-optimal probabilistic RNA-seq quantification. Nat Biotechnol 34, 525–527, doi:10.1038/nbt.3519 (2016).

41 R: A Language and Environment for Statistical Computing} v. 4.1.0 (R Foundation for Statistical Computing, Vienna, Austria, 2021).

42 gplots: Various R Programming Tools for Plotting Data v. 3.0.1.1 (2019).

43 Everitt, B. S. L., Sabine; Leese, Morven; Stahl, Daniel. Cluster Analysis, 5th Edition. (Wiley, 2011).

44 Tibshirani, R., Walther, G. & Hastie, T. Estimating the number of clusters in a data set via the gap statistic. J. R. Statist. Soc. *B* 63, 411±423 (2001).

45 Charrad, M., Ghazzali, N., Boiteau, V. & Niknafs, A. NbClust: An {R} Package for Determining the Relevant Number of Clusters in a Data Set}. Journal of Statistical Software 61, 1-36 (2014).

46 Torgerson, W. S. Multidimensional scaling, I: theory and method. Psychometrika 17, 401–419 (1952).

47 Wilcoxon, F. Individual comparisons by ranking methods. Biometrics Bulletin 1, 80–83 (1945).

48 Benjamini, Y. & Hochberg, Y. Controlling the false discovery rate; A practical and powerful approach to multiple testing. J. Roy. Stat. Soc. Ser. B 57, 289–300 (1995).

