## Supplementary Materials for "A pancreatic cancer mouse model with human immunity"

Fig. S1. HLA typing of PANC-1 cells.

Fig. S2. Macroscopic images of 7-week-old orthotopic PANC-1 tumors grown in nude mice.

Fig. S3. Profile of HIS-derived splenocytes used for *in vitro* co-culture studies

Fig. S4. The effect of PDAC cells on the maintenance of HLA-matched immune cells

Fig. S5. HLA typing of 0799E cells

Fig. S6. Macroscopic images of 7-week-old orthotopic 0799E tumors grown in nude mice

Fig. S7. Representative images of an orthotopic tumor and the liver harvested from 0799E huPDAC mice

Fig. S8. Hierarchical clustering of PDAC-infiltrating immune cells based on CIBERSORTx analysis

Fig. S9. PDAC-resident Tregs express the αvβ5 integrin.

Fig. S10. Representative flow cytometry analysis of CD8/Treg ratio in the tumor from PANC-1 huPDAC mice treated with or without iRGD.

Fig. S11. The effect of iRGD treatment on the growth of PANC-1 huPDAC tumors

Fig. S12. Gross appearance of the liver of PANC-1 huPDAC mice treated with iRGD

Fig. S13. A representative high magnification H&E image of a liver section prepared from iRGD-treated PANC-1 huPDAC mice

Table. S1. Overview of the treatment performed in PANC-1 huPDAC mice

Table. S2. Statistical analysis of the proportion of immune cell fractions between huPDAC tumors and TCGA patient-derived PDAC samples.

Table. S3. Statistical analysis of the change in proportion of immune cell fractions iRGD treated vs huPDAC tumors a

| Typing results with NMDP code | | | | | | |
|---|---|---|---|---|---|---|
| Sample Name | A1 | A2 | B1 | B2 | DRB11 | DRB12 |
| PANC-1 | 02:CEJST | 11:CMVFM | 38:CEJVS | 38:CEJVS | 13:BZEMT | 13:BZEMT |

b

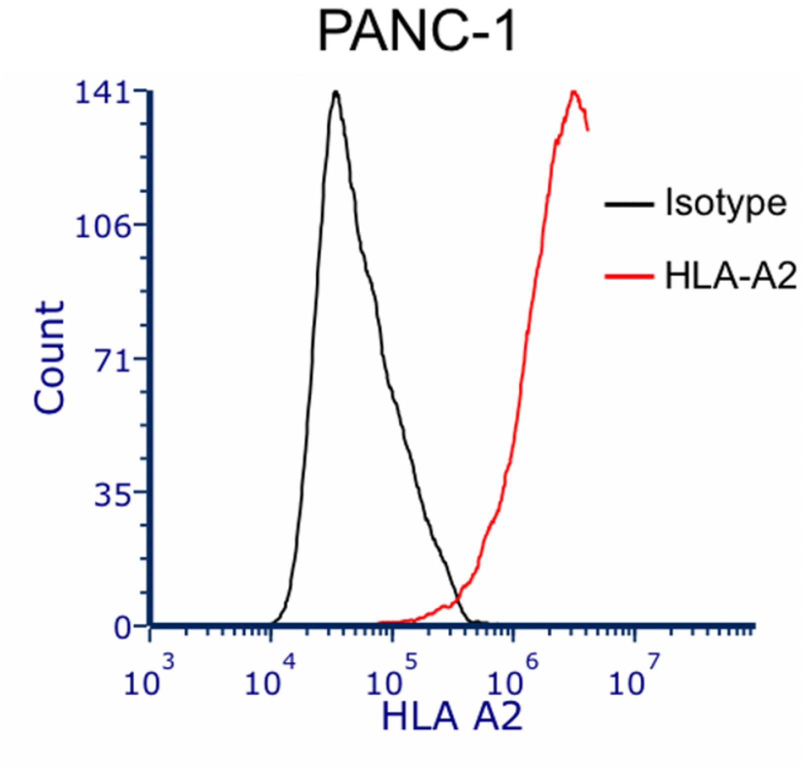


**Supplementary Figure 1. HLA typing of PANC-1 cells.** (a) HLA-A, -B and -DRB1 alleles are shown by National Marrow Donor Program (NMDP) code. (b) Expression of HLA-A2 on PANC-1 cells analyzed by flow cytometry. Black, isotype control; red, HLA-A2 antibody.

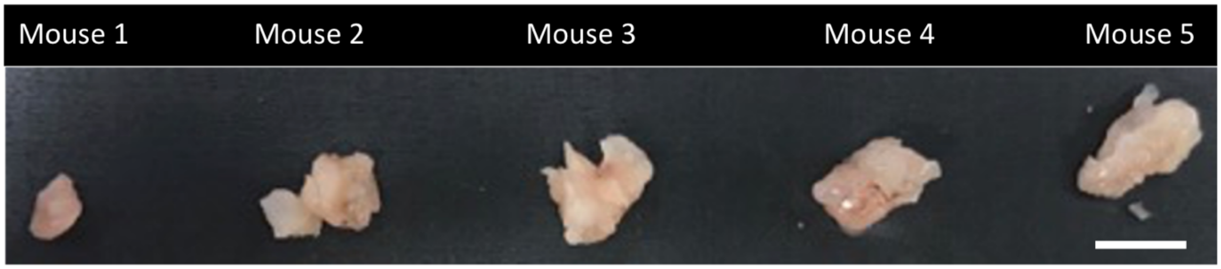


**Supplementary Figure 2. Macroscopic images of 7-week-old orthotopic PANC-1 tumors grown in nude mice.** Scale bar, 10 mm.

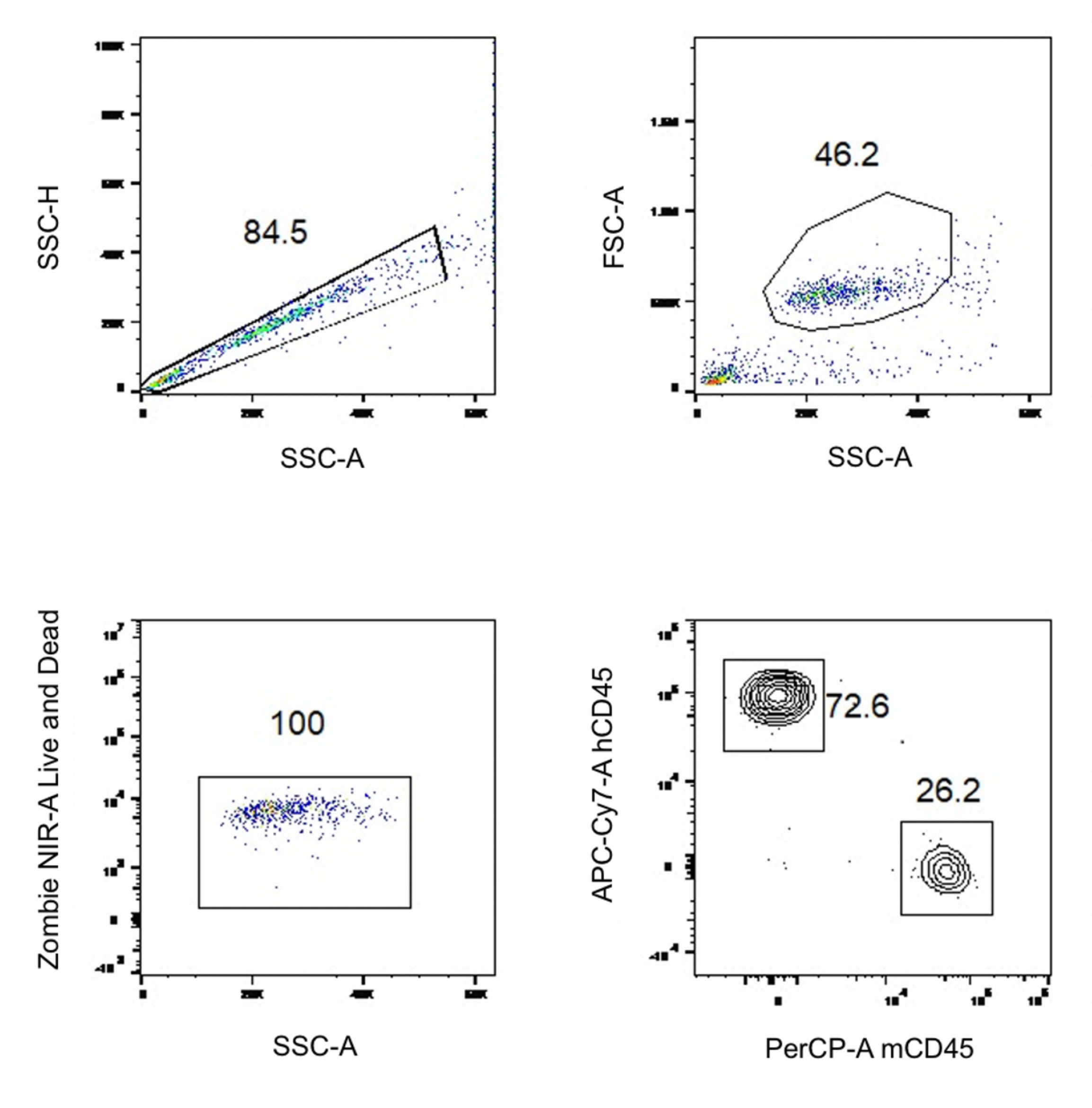


**Supplementary Figure 3. Profile of HIS-derived splenocytes used for *in vitro* co-culture studies.** Splenocytes harvested from HIS mice were subjected to flow cytometry. A side scatter (SSC)-A/SSC-H plot was used to gate single cell populations, and a forward scatter (FSC)-A/SSC-A plot was used to gate live cells. Zombie was used to confirm the presence of live cells, and the proportion of human ($hCD45^+$) and mouse $CD45^+$ ($mCD45^+$) cells was studied.

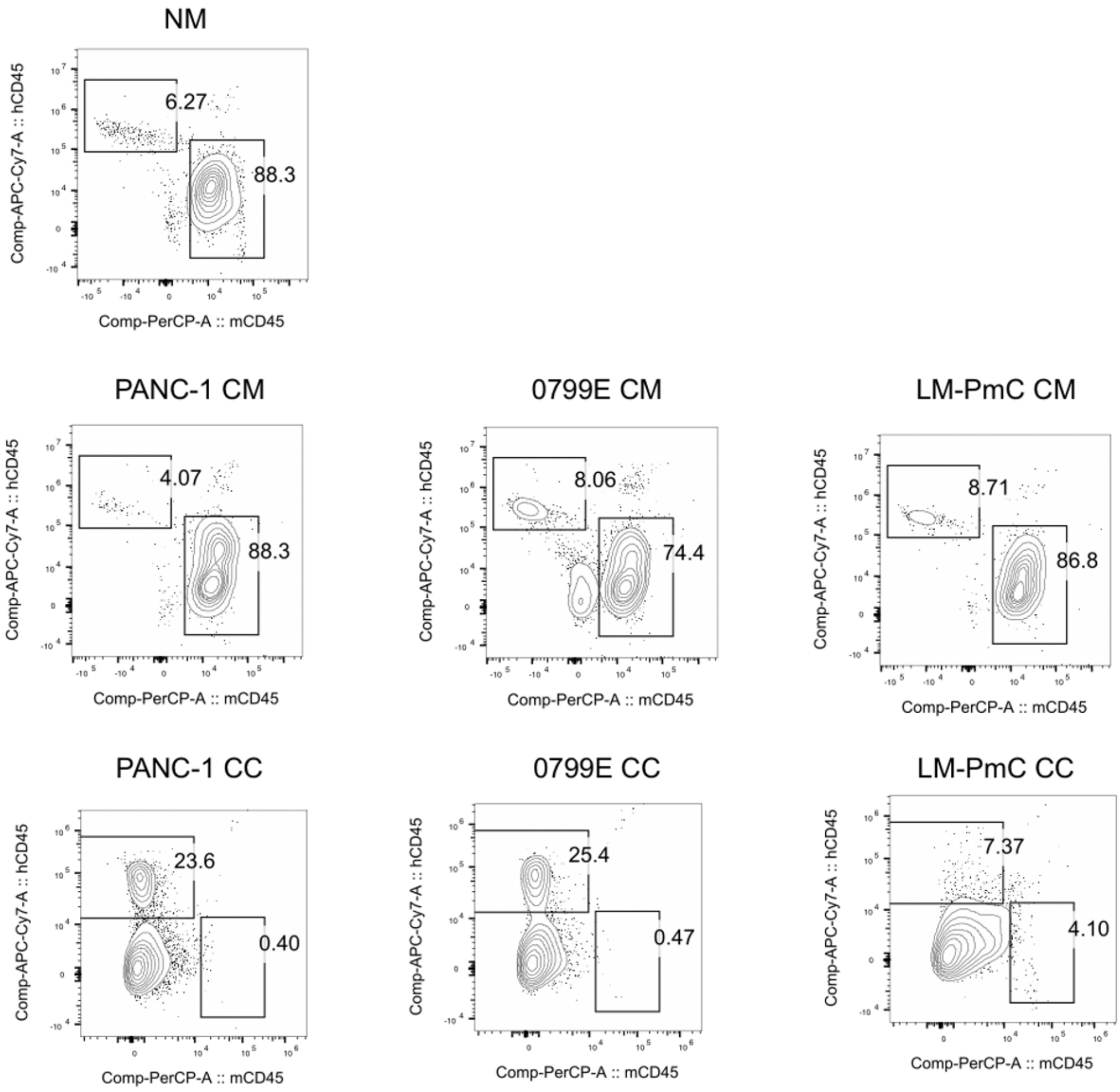


**Supplementary Figure 4. The effect of PDAC cells on the maintenance of HLA-matched immune cells.** Splenocytes isolated from the spleen of HIS mice were cultured in normal media (NM; 10% FBS containing DMEM), in conditioned media (CM) prepared from 3-day-old cultures of PANC-1) or on top of a 3-day-old monolayer prepared with PANC-1, 0799E, or LM-PmC cells. Proportion of $hCD45^{+}$ and $mCD45^{+}$ cells were analyzed by flow cytometry.

a

| Typing results with NMDP code | | | | | | |
|---|---|---|---|---|---|---|
| Sample Name | A1 | A2 | B1 | B2 | DRB11 | DRB12 |
| 0799E | 02:CEJST | 02:CEJST | 51:CMVGS | 51:CMVGS | 15:CEKDG | 15:CEKDG |

b

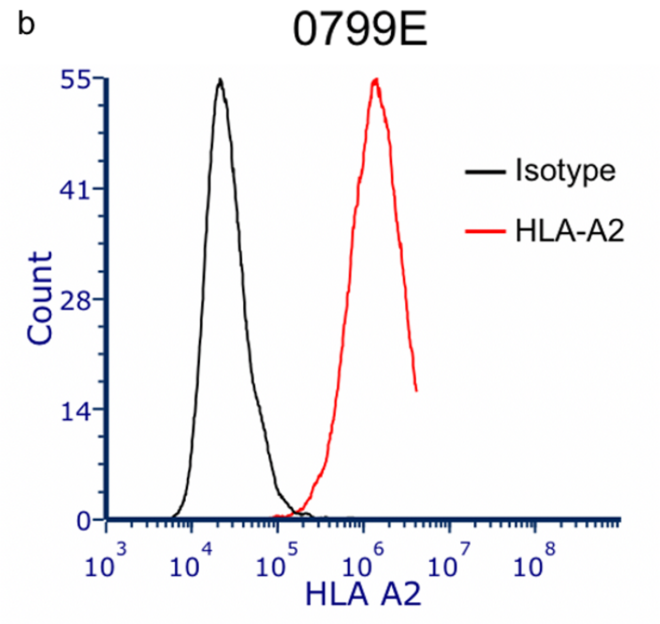


**Supplementary Figure 5. HLA typing of 0799E cells.** (a) HLA-A, -B and -DRB1 alleles are shown using the NMDP code. (b) Expression of HLA-A2 on 0799E cells analyzed by flow cytometry. Black, isotype control; red, HLA-A2 antibody.

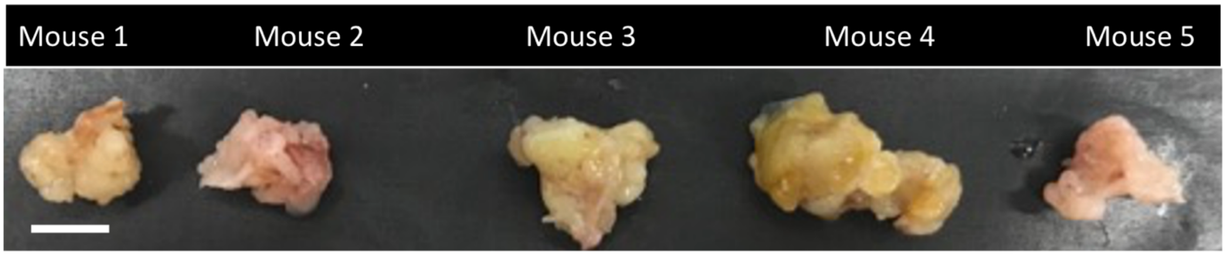


**Supplementary Figure 6. Macroscopic images of 7-week-old orthotopic 0799E tumors grown in nude mice.** Scale bar, 10 mm.

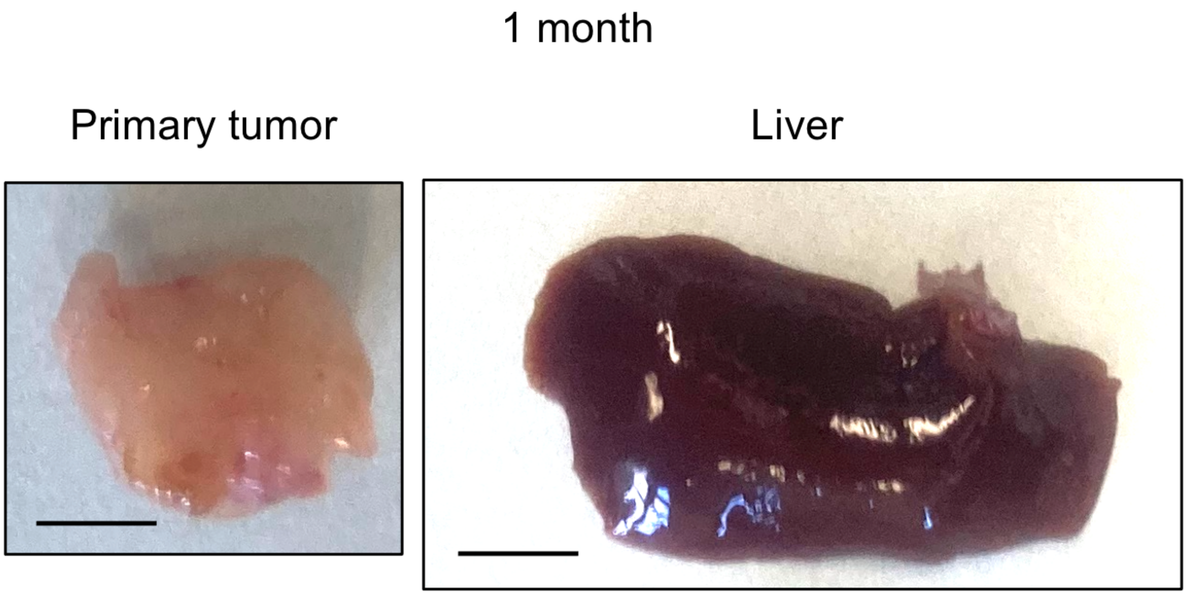


**Supplementary Figure 7. Representative images of an orthotopic tumor and the liver harvested from 0799E huPDAC mice.** The specimens were harvested 1 month after tumor implantation. Scale bar, 5 mm.

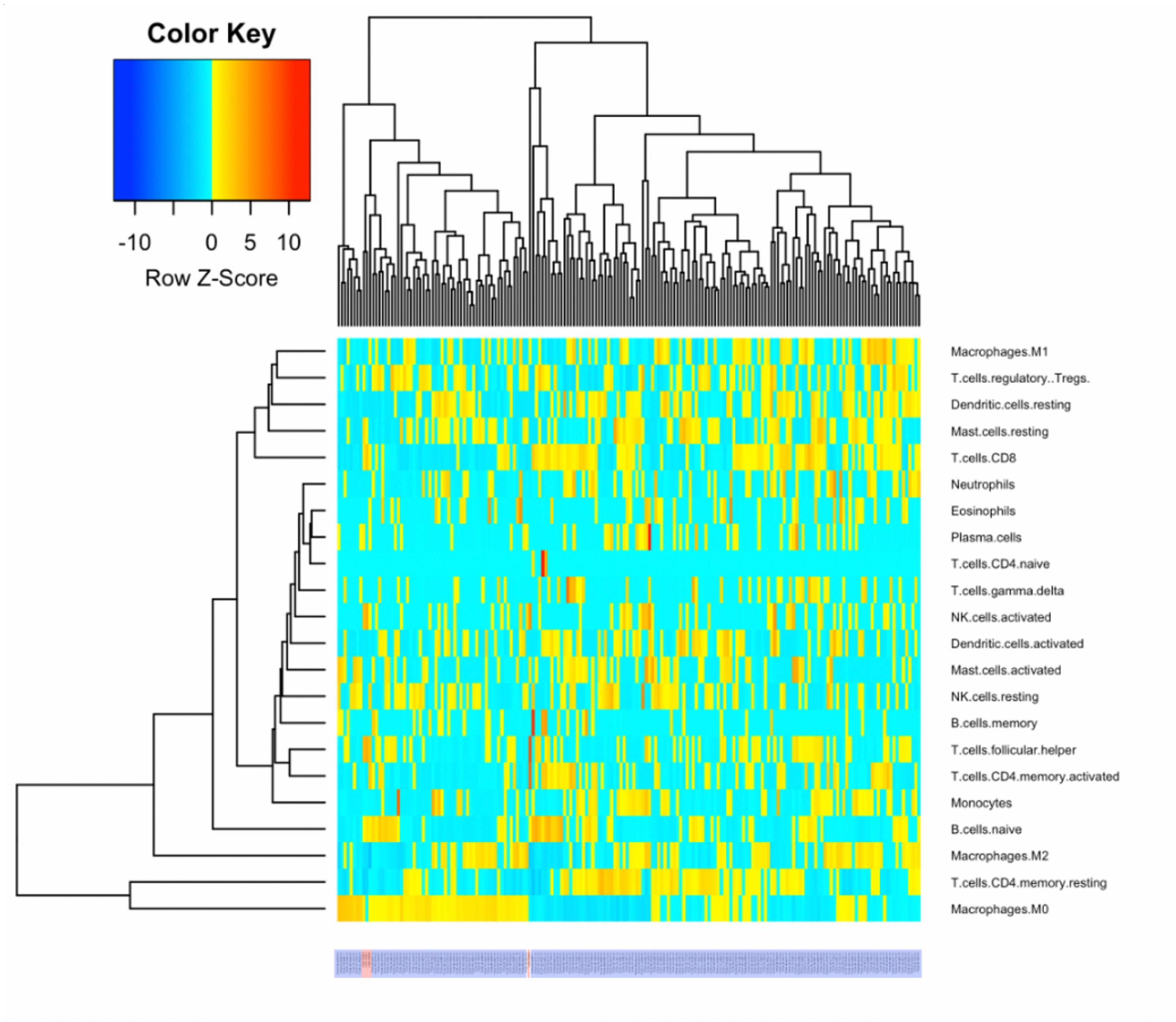


**Supplementary Figure 8. Hierarchical clustering of PDAC-infiltrating immune cells based on CIBERSORTx analysis.** Hierarchical clustering of major immune cell fractions in 2 of PANC-1 or 2 of 0779E huPDAC tumors (samples coded in red at the bottom) and 182 patient-derived PDAC samples in the TCGA database (samples coded in blue at the bottom) was performed based on CIBERSORTx analysis of the RNA-seq data. A dendrogram and heatmap are shown. Calculations were performed and images generated using the heatmap.2 command in gplots.

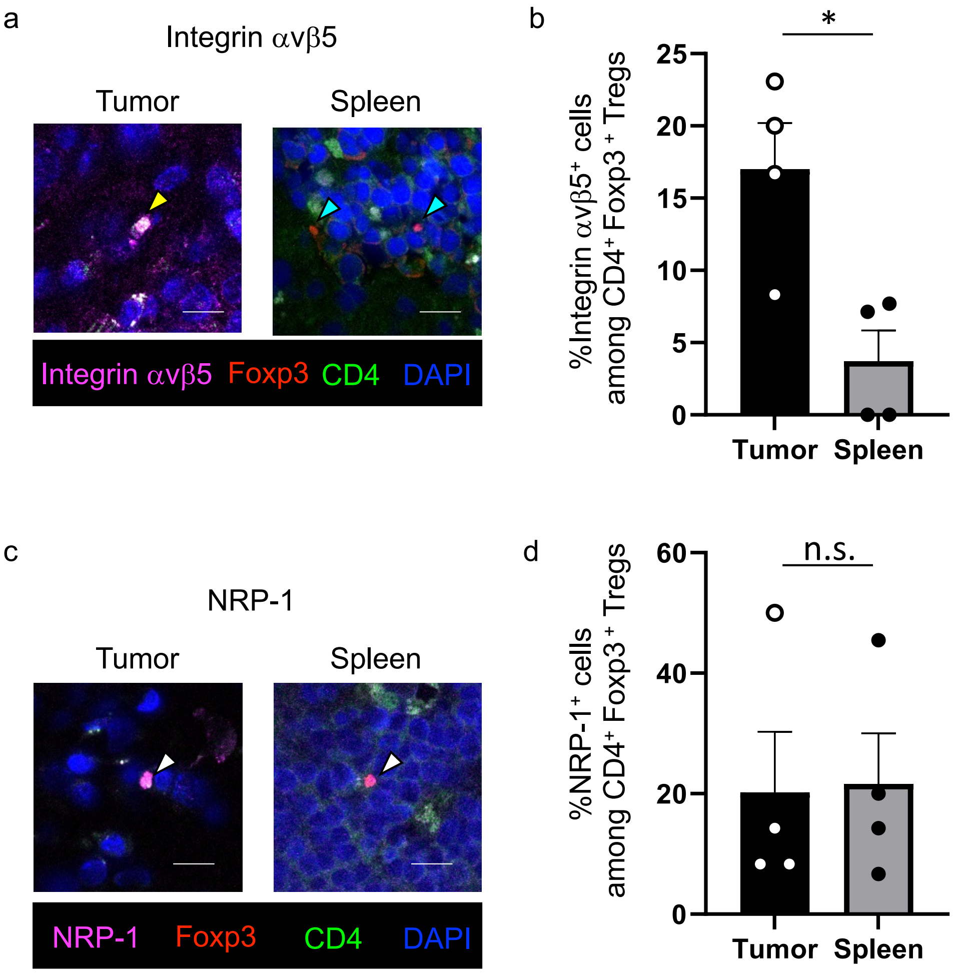


**Supplementary Figure 9. PDAC-resident human Tregs express the αvβ5 integrin.** (a, c) Representative confocal images of αvβ5 integrin$^{+}$ Foxp3$^{+}$ CD4$^{+}$ human Tregs (a) or NRP-1$^{+}$ Foxp3$^{+}$ CD4$^{+}$ human Tregs (c) in the tumor and spleen in huPDAC mice. Magenta, αvβ5 integrin (a) or NRP-1 (c); red, Foxp3; green, CD4; blue, DAPI. Yellow arrow head, αvβ5 integrin$^{+}$ Foxp3$^{+}$ CD4$^{+}$ Tregs; cyan arrow heads, αvβ5 integrin$^{neg}$ Foxp3$^{+}$ CD4$^{+}$ Tregs; white arrow heads, NRP-1$^{+}$ Foxp3$^{+}$ CD4$^{+}$ Tregs. (b, d) Proportion of αvβ5 integrin-positive (b) or NRP-1-positive (d) cells among Foxp3$^{+}$ CD4$^{+}$ Tregs in the tumor and spleen based on (a and c). n = 4. Error bars, mean ± standard error: Student's t test; *$p < 0.05$; n.s., not significant.

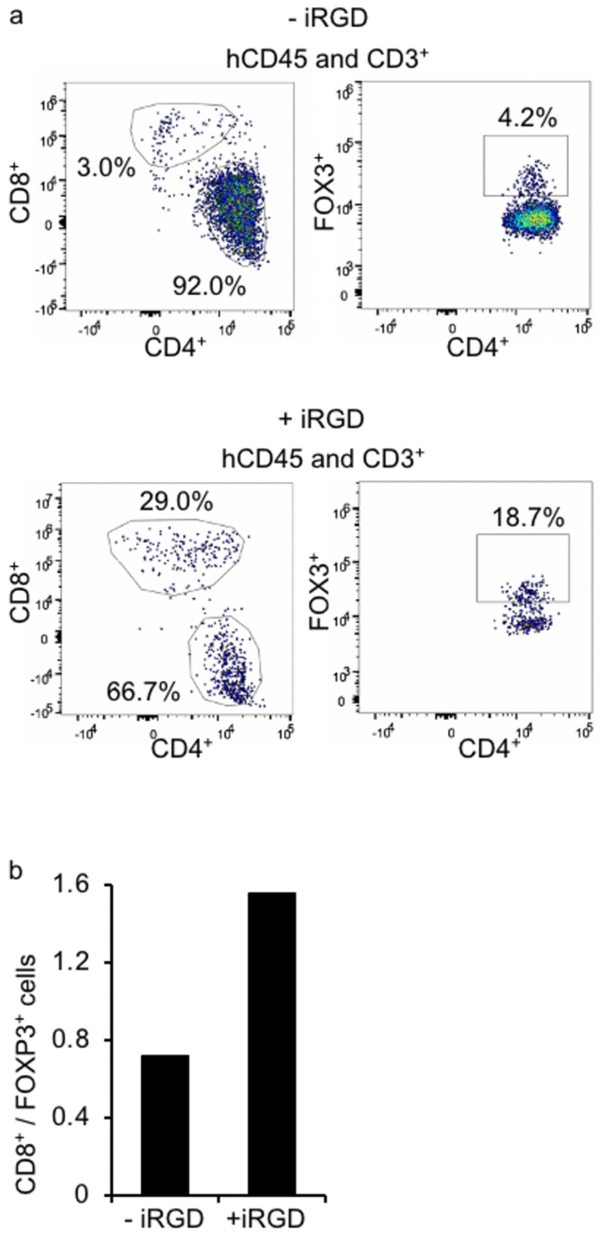


**Supplementary Figure 10. An example of flow cytometry analysis of CD8/Treg ratio in the tumor from PANC-1 huPDAC mice treated with or without iRGD.** (a) Proportion of $CD4^+$, $CD8^+$, and $FOXP3^+$ T cells among $hCD45^+$ and $CD3^+$ cells in PANC-1 huPDAC tumors that received treatment with or without iRGD. (b) The CD8/Treg ratio was calculated based on (a).

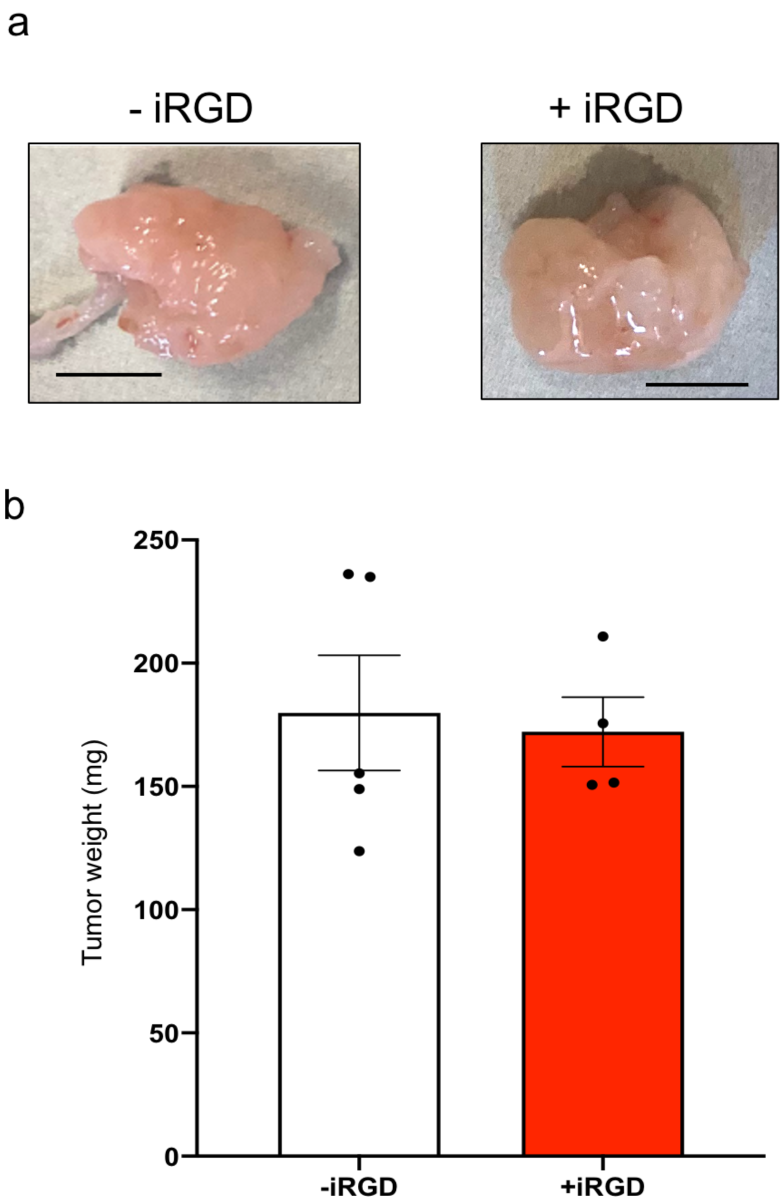


**Supplementary Figure 11. The effect of iRGD treatment on the growth of PANC-1 huPDAC tumors.** huPDAC mice bearing orthotopic PANC-1 tumors were treated with (n = 4) or without (n = 5) 1.2 mg / kg of iRGD 3 times a week for 8 weeks. (a) A representative macroscopic appearance of the PDAC. (b) The wet weight of the orthotopic tumors was measured at the end of the treatment.

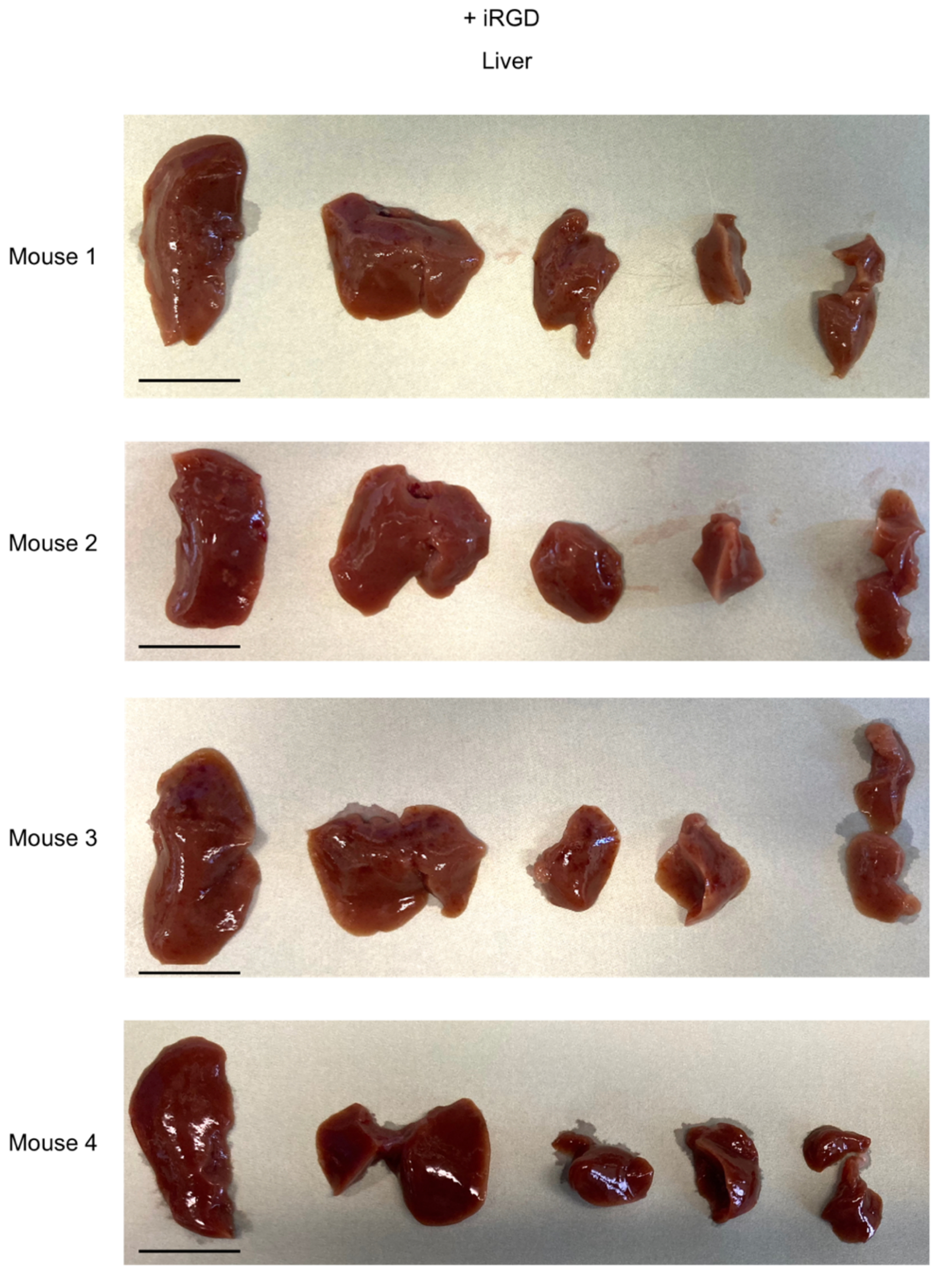


**Supplementary Figure 12. Gross appearance of the liver of PANC-1 huPDAC mice treated with iRGD.** The liver was harvested from orthotopic PANC-1 huPDAC mice treated 1.2 mg / kg of iRGD 3 times a week for 8 weeks (n = 4). The liver was divided into multiple segments to show the entire tissue. Note that there are no visible tumor nodules on the liver surface. Scale bar, 10 mm.

+ iRGD

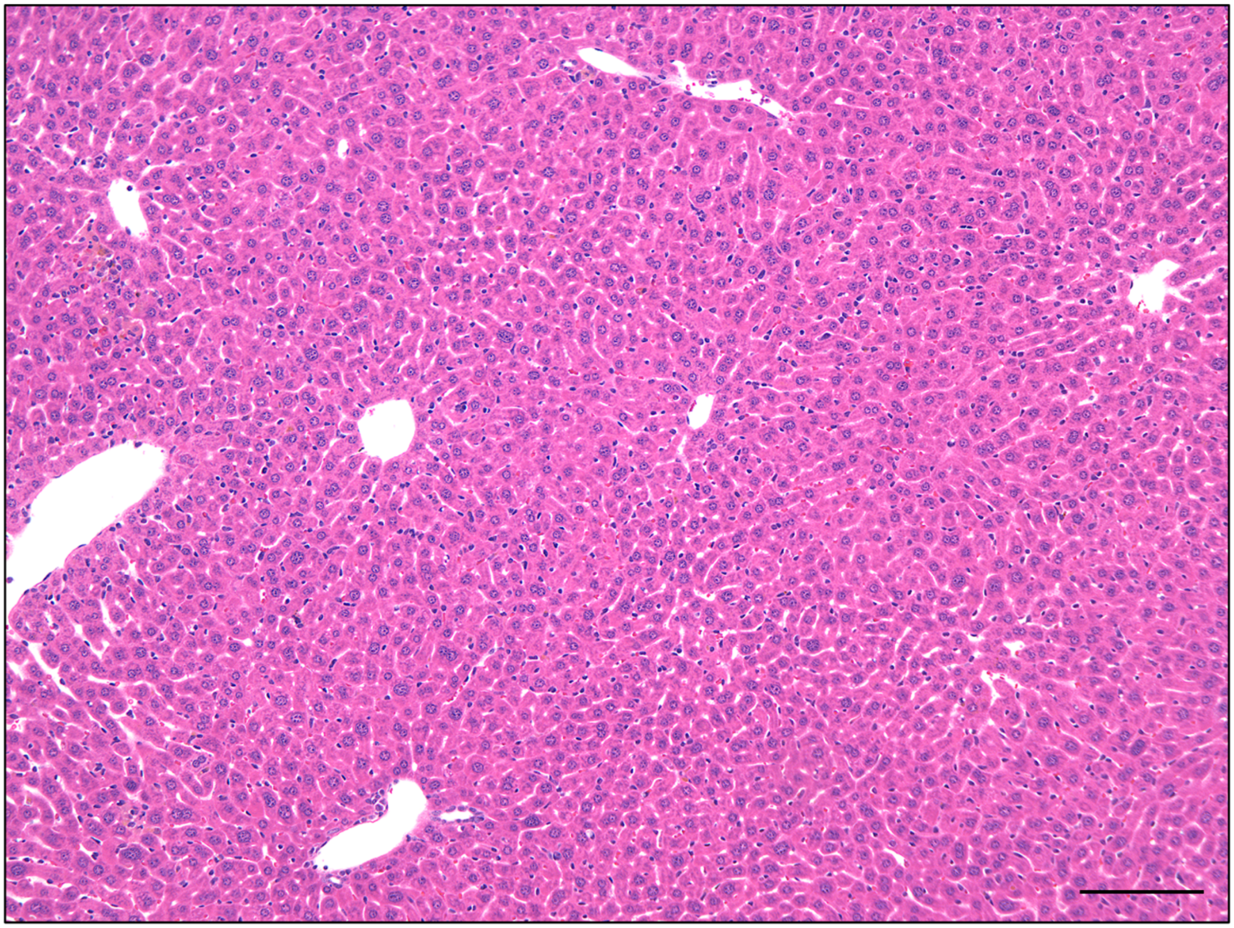

**Supplementary Figure 13. A representative high magnification H&E image of a liver section prepared from iRGD treated huPDAC mice.** The liver samples shown in Supplementary Fig. 11 were sectioned, stained with H&E, and subjected to microscopy. Scale bar, 100 μm

Supplementary Table. 1. Overview of the treatment performed in PANC-1 huPDAC mice

| No. | $hCD45^+$ (%) before Transplantation | Tolerance | CD8 depletion | iRGD | $hCD45^+$ (%) | | | |
|---|---|---|---|---|---|---|---|---|
| | | | | | Tumor | Metastasis | Spleen | PBMC |
| 1 | 94.8 | YES | YES | NO | 85.9 | NA | 69.0 | NA |
| 2 | 93.2 | YES | NO | NO | 85.3 | 90.2 | 72.1 | NA |
| 3 | 34.4 | NO | NO | NO | 88.9 | 88.8 | 17.7 | 5.9 |
| 4 | 21.5 | NO | NO | NO | 91.5 | 95.4 | 14.5 | 5.9 |
| 5 | 8.0 | NO | NO | NO | 96.0 | 64.1 | 31.9 | 6.7 |
| 6 | 77.0 | NO | NO | YES | 78.4 | | 82.3 | 63.6 |
| 7 | 65.9 | NO | NO | YES | ND | | 75.2 | 57.1 |
| 8 | 83.9 | NO | NO | YES | 85.0 | | 78.5 | 46.0 |
| 9 | 12.7 | NO | NO | YES | 48.0 | | 6.1 | 0.5 |

NA, Not applicable; ND, Not detected.

**Supplementary Table. 2. Statistical analysis of the proportion of immune cell fractions between huPDAC tumors and TCGA patient-derived PDAC samples.** False discovery rate (Fdr) < 0.05 was considered statistically significant.

| | deltamean | p.value | fdr |
|---|---|---|---|
| NK cells, activated | 0.055 | 0.001 | 0.017 |
| Follicular helper T | 0.134 | 0.003 | 0.027 |
| CD4 memory T, resting | -0.101 | 0.004 | 0.027 |
| Macrophages, M2 | -0.097 | 0.005 | 0.027 |
| DC, resting | -0.041 | 0.011 | 0.047 |
| Neutrophils | -0.009 | 0.025 | 0.090 |
| DC, activated | -0.016 | 0.043 | 0.134 |
| NK cells, resting | -0.003 | 0.054 | 0.148 |
| B cells, naïve | 0.036 | 0.089 | 0.217 |
| T cells, gamma delta | -0.008 | 0.163 | 0.358 |
| CD8 T | 0.035 | 0.270 | 0.540 |
| B cells, memory | -0.006 | 0.300 | 0.551 |
| Plasma cells | 0.000 | 0.387 | 0.655 |
| Mast cells, activated | -0.002 | 0.488 | 0.739 |
| Macrophages, M1 | -0.005 | 0.504 | 0.739 |
| Tregs | -0.016 | 0.598 | 0.811 |
| CD4 memory T, activated | 0.050 | 0.626 | 0.811 |
| Monocytes | -0.009 | 0.734 | 0.847 |
| Mast cells, resting | 0.001 | 0.740 | 0.847 |
| Eosinophils | -0.002 | 0.770 | 0.847 |
| CD4 naïve T | -0.001 | 0.812 | 0.851 |
| Macrophages, M0 | 0.005 | 0.883 | 0.883 |

**Supplementary Table. 3. Statistical analysis of the change in proportion of immune cell fractions iRGD treated vs huPDAC tumors.** False discovery rate (Fdr) < 0.05 was considered statistically significant. NA, Not applicable

| | deltamean | p.value | fdr |
|---|---|---|---|
| B cells, naïve | 0.050 | 0.057 | 0.314 |
| CD8 T | -0.147 | 0.057 | 0.314 |
| Monocytes | -0.014 | 0.057 | 0.314 |
| Macrophages, M2 | -0.065 | 0.057 | 0.314 |
| Macrophages, M0 | 0.082 | 0.114 | 0.435 |
| Plasma cells | 0.003 | 0.119 | 0.435 |
| CD4 memory T, resting | 0.071 | 0.229 | 0.629 |
| Macrophages, M1 | 0.022 | 0.229 | 0.629 |
| Eosinophils | 0.002 | 0.270 | 0.661 |
| CD4 memory T, activated | 0.055 | 0.359 | 0.733 |
| Tregs | -0.030 | 0.400 | 0.733 |
| Mast cells, resting | -0.041 | 0.400 | 0.733 |
| Mast cells, activated | 0.004 | 0.582 | 0.985 |
| DC, resting | -0.001 | 0.696 | 1 |
| NK cells, resting | -0.003 | 0.854 | 1 |
| DC, activated | -0.005 | 0.854 | 1 |
| Follicular helper T | 0.023 | 0.857 | 1 |
| NK cells, activated | -0.007 | 0.857 | 1 |
| B cells, memory | 0.000 | NA | NA |
| CD4 naïve T | 0.000 | NA | NA |
| T cells, gamma delta | 0.000 | NA | NA |
| Neutrophils | 0.000 | NA | NA |
